# The pro-inflammatory response to influenza A virus infection is fueled by endothelial cells

**DOI:** 10.1101/2022.08.19.504520

**Authors:** Lisa Bauer, Laurine C. Rijsbergen, Lonneke Leijten, Feline F. W. Benavides, Danny Noack, Mart M. Lamers, Bart. L. Haagmans, Rory D. de Vries, Rik L. de Swart, Debby van Riel

**Affiliations:** Department of Viroscience, Erasmus MC, Rotterdam, the Netherlands

**Keywords:** endothelial cells, inflammation, influenza A virus, airway organoids

## Abstract

Morbidity and mortality from influenza are associated with high levels of systemic inflammation. Endothelial cells have been shown to play a key role in this systemic inflammatory response during severe influenza A virus (IAV) infections, despite the fact that these are rarely infected in humans. However, how endothelial cells contribute to these systemic inflammatory responses is unclear. To investigate this, we developed a transwell-system in which airway organoid-derived differentiated human lung epithelial cells at the apical side were co-cultured with primary human lung microvascular endothelial cells (LMEC) at the basolateral side. We compared the susceptibility of endothelial cells to pandemic H1N1 virus isolated in 2009 and seasonal H1N1 and H3N2 virus isolated in 2019, and assessed the associated immune responses. Despite the detection of IAV nucleoprotein in LMEC monocultures, there was no evidence for productive infection. In epithelial-endothelial co-cultures, abundant IAV infection of epithelial cells resulted in the breakdown of the epithelial barrier, but infection of LMECs was rarely detected. Furthermore, we observed a significantly higher secretion of pro-inflammatory cytokines in LMECs when co-cultured with IAV-infected epithelial cells, compared to LMEC monocultures exposed to IAV. Taken together, our data show that endothelial cells are abortively infected by IAV, but can fuel the inflammatory response. As endothelial cells are a prominent cell type in the lung, it is possible that they play an important role in the systemic inflammatory response during IAV infections.

## Introduction

Influenza A virus (IAV) infections often cause mild, self-limiting respiratory disease, but can also result in severe disease with high morbidity and mortality. Severe disease occurs especially in risk groups such as young infants, pregnant women, patients with comorbidities and the geriatric population ^1,2^. Severe IAV infections are not exclusively induced by virus replication itself, but also by the host immune response ^1,3,4^. For example, high levels of pro-inflammatory responses detected in the blood, are associated with high morbidity in pandemic in H1N1 viruses, either from 1918 or 2009. and zoonotic IAV infections in humans, or in nonhuman primates ^5–7^. This inflammatory response is often referred to as a cytokine storm ^7–9^.

Endothelial cells are involved in the early cytokine and chemokine response as well as the recruitment of innate immune cells to the lung during an IAV infection in mammals. Suppression of early innate immune responses in endothelial cells and associated cytokine production greatly reduced the mortality of experimentally infected animals ^10^. However, how endothelial cells contribute to the systemic immune responses is unclear. IAV antigen is rarely detected in human endothelial cells *in vivo* after infection with zoonotic, pandemic or seasonal IAV in humans or animal models. ^10–12^. However, *in vitro* several studies show that IAV with multibasic cleavage sites such as H5N1 and H7N9 productively replicate in human endothelial cells ^13–17^. In contrast, IAV without multibasic cleavage sites such as H1N1, H2N2 and H3N2 virus cause abortive infections of endothelial cells but induce a wide array for pro-inflammatory cytokines ^10,11^. Therefore, it is hypothesized that seasonal or pandemic IAV might augment a pro-inflammatory response of endothelial cells through intracellular pathogen recognition receptors (PRR) sensing viral RNA ^18–21^. Cytokines and chemokines produced by endothelial cells would directly be released into the blood stream and contribute to the systemic inflammatory response^20,21^.

To investigate the contribution of endothelial cells to the pro-inflammatory response during IAV infection, we developed an *in vitro* trans-well model in which epithelial cells (airway organoids cultured at air-liquid-interface (AO at ALI) and endothelial cells (primary lung microvascular endothelial cells (LMEC)) are co-cultured, to mimic the *in vivo* situation in the lower respiratory tract. This model allowed us to investigate the contribution of endothelial cells to pro-inflammatory cytokine production after infection of epithelial cells with pandemic or seasonal IAV.

## Results

### Inefficient replication of influenza A virus in lung microvascular endothelial cells elicits an interferon response

To assess the susceptibility and permissiveness of primary LMECs to IAV infection, we cultured LMECs either at the apical or basolateral side of a filter in a transwell system and subsequently inoculated cells from the apical side with pandemic H1N1 2009 virus (pH1N1), or seasonal H1N1 or H3N2 viruses isolated in 2019. In LMECs cultured at the apical or basolateral side no increase of infectious virus titers was detected in the apical or basolateral compartments (Figure 1A, B). Virus nucleoprotein (NP) antigen was detected in some virus inoculated LMECs 24 hours post-inoculation (hpi) by immunofluorescence (IF) staining in both experimental set-ups (Figure 1C, D). Flow cytometry analysis revealed 1-2% IAV infected cells at 24 hpi and <1% at 72 hpi of NP^+^ LMECs. pH1N1 virus infected significantly more LMECs compared to H1N1 and H3N2 virus at 24 (∼2% versus ∼1%) and 72 hpi (∼1% versus 0.3%) (Figure 1E). To confirm infection of LMECS, we quantified viral RNA genome copies by quantitative real-time PCR (qRT-PCR) over time, and showed that intracellular viral genome replication peaked at 24 hpi for all three viruses, after which the amount of genome copies declined (Figure 1F).

**Figure 1.**
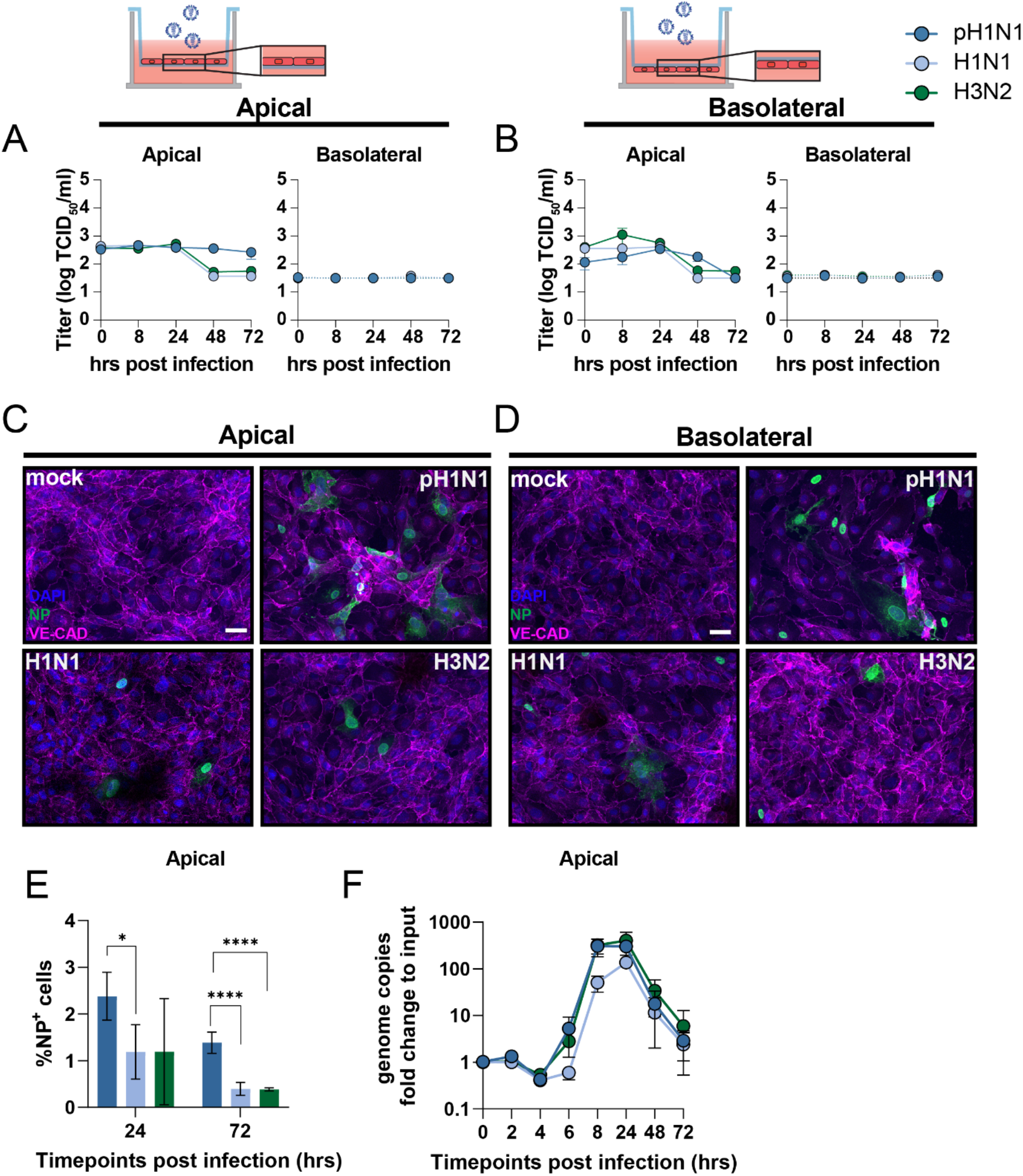
Abortive infection of influenza A virus in lung microvascular endothelial cells. To evaluate replication efficiency, lung microvascular endothelial cells (LMECs) were plated on the (A) apical or (B) basolateral side of a transwell filter. LMECs were inoculated with pH1N1, H1N1, or H3N2 virus at MOI 1 and at the indicated time points supernatants of the apical as well as basolateral side were harvested and virus titers were determined by endpoint titration. Infection efficiency was determined by immunofluorescence staining. LMECs plated on the (C) apical and (D) basolateral were inoculated with pH1N1, H1N1, or H3N2 virus. Cells were fixed 24 hours post inoculation and stained for the endothelial cell marker Vascular Endothelial-Cadherin (VE-CAD, magenta) and influenza A virus nucleoprotein (NP (green). Hoechst (blue) was used to visualize nuclei. (E) Percentage of infection determined by flow cytometry at 24 and 72 hours post inoculation. (F) Viral RNA genome copies were quantified by quantitative real time PCR at indicated time points. Data represent mean +/- standard deviation (SD) from at least three independent experiments performed in biological duplicates and flow cytometry was performed in biological triplicates. A one-way ANOVA multiple comparison test was used to compare groups (*< 0.05, **<0.01, ***<0.005). Scale bar: 20 μm.

To evaluate whether infection of LMECs induced an antiviral immune response, we inoculated LMECs with pH1N1, H1N1 or H3N2 virus and measured *interferon-β* (*IFN-β), IFN-λ* and interferon-stimulated gene (ISG) *IFIT1* 24 hpi (Supplementary figure 1). As a positive control, we exposed LMECs either to recombinant IFN-β, IFN-λ, or to Toll-like receptor 3 agonist Polyinosinic:polycytidylic acid (Poly I:C), mimicking double stranded RNA, either in the supernatant or via transfection. Upon pH1N1, H1N1 or H3N2 virus infection *IFN-β, IFN-λ* and *IFIT1* were significantly upregulated as compared to the mock control. In the positive control conditions *IFN-β, IFN-λ* and *IFIT1* were significantly induced, except after IFN-λ treatment where only *IFIT1* and *IFN-λ* were significantly induced. This suggests that LMECs express interferon α/β receptor (IFNAR) but not IFNLR. In conclusion, there is intracellular viral genome replication of pH1N1, H1N1 and H3N2 viruses in LMECs, however we do not detect infectious virus progeny in the supernatant, which is a sign of abortive infection. The intracellular genome replication induces a type-I and type-III interferon response.

### Replication of influenza A viruses in epithelial cell single cultures and epithelial-endothelial cell co-cultures induces a robust cytokine response

Epithelial cells are the first cells to be infected by IAV in the lung. We used AO at ALI as our source of respiratory epithelial cells and validated that this was a pseudostratified layer containing ciliated and goblet cells (Supplemental figure 2A and 2B, respectively). Next, we developed a co-culture model using transwells containing AO at ALI at the apical side and LMECs at the basolateral side of the transwell Pandemic H1N1, H1N1 or H3N2 virus inoculation of epithelial cells resulted in efficient virus replication in the single epithelial cultures as well as co-cultures measured in the apical compartment (Supplemental figure 2C and Figure 2A). All viruses replicated slightly faster, measured by viral titers in the apical compartment, in co-cultures compared to the mono-cultures, but to ∼1log lower virus titers (Supplementary figure 3A). In the basolateral compartment infectious virus was only detected at 48 hpi in the single cultures, which was associated with a reduction in the transepithelial electrical barrier (TER) (Supplemental figure 2C-D). No infectious virus particles were detected in the basolateral compartment of the co-cultures (Figure 2A). IAV infected ciliated epithelial cells could be detected at 24 hpi with minimal visible cell damage using a hematoxylin and eosin staining (Figure 2B). At 72 hpi flattening of the epithelial cell layer was observed associated with hypertrophic epithelial cells and loss of ciliated epithelial cells. Loss of cilia was confirmed using an IF staining for acetylated α-tubulin and there were clearly less cilia at 72 hpi. In addition, we observed loss of the tight junction marker zona-occludin 1. In IAV infected LMECS we detected changes in the expression of vascular endothelial cadherin, which was more diffuse compared to uninfected cells where it was predominantly observed on the cell surface. In the LMECs there were also some structural changes visualized by vascular endothelium cadherin (Figure 2C-D).

**Figure 2.**
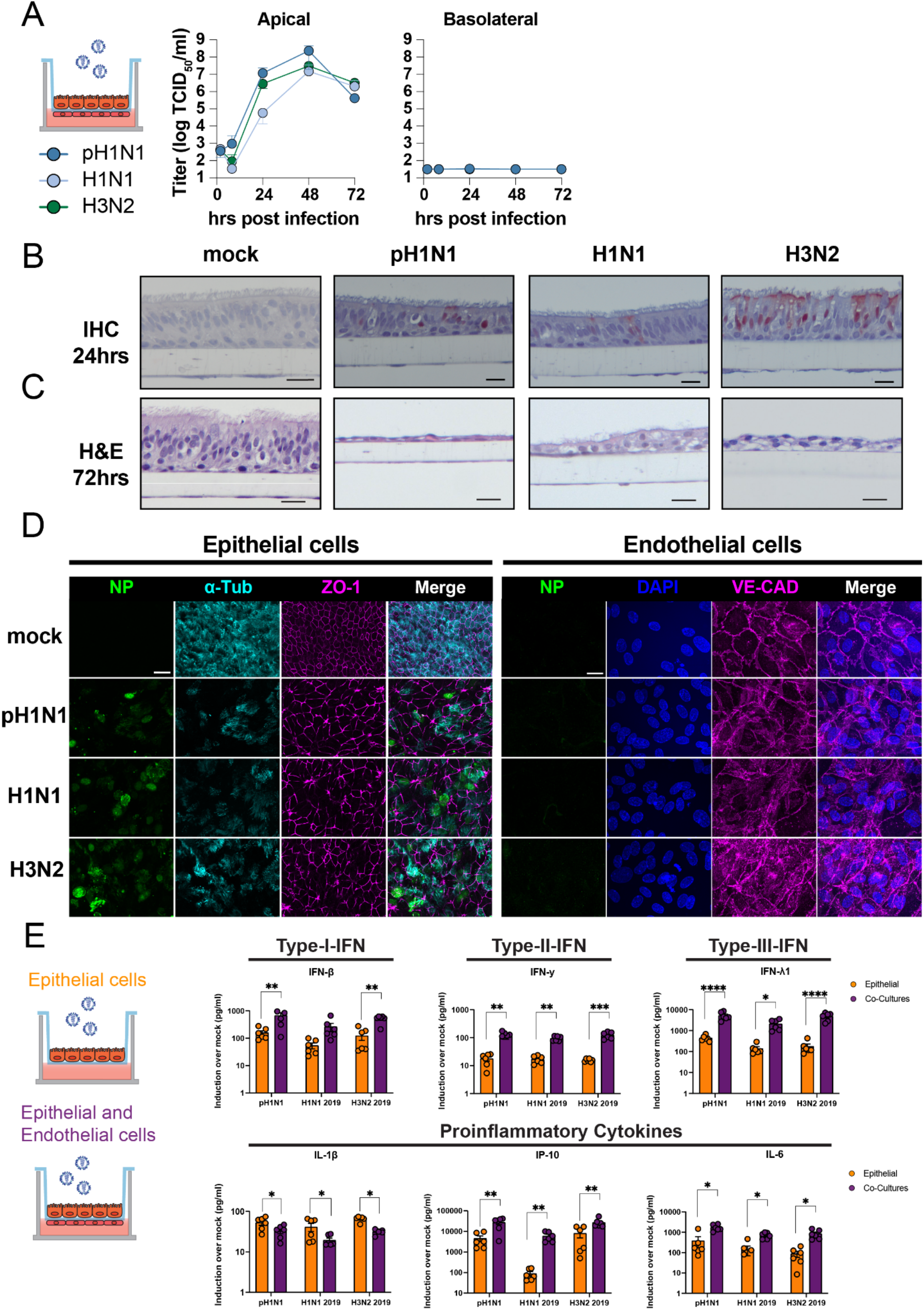
Influenza A virus infection and cytokine profiling in epithelial cell cultures alone of in co-culture with endothelial cells. (A) Well-differentiated airway organoids at air-liquid interface (AO at ALI) in co-culture with lung microvascular endothelial cells (LMECs) were inoculated with pH1N1, H1N1 or H3N2 virus at MOI 1. At the indicated timepoints virus titers were determined in the supernatants of the apical and basolateral compartments. (B) Detection of influenza A virus (IAV) nucleoprotein (NP) by immunohistochemistry of the AO at ALI-LMEC co-cultures 24 hours post inoculation (C) Hematoxylin and eosin (H&E) staining of the co-cultures 72 hours post inoculation (scale bar 20 μm) (D) At 72 hours post inoculation well-differentiated AO at ALI were stained for IAV NP (green), the cilia marker acetylated-α-tubulin (cyan) and the tight-junction marker Zona occludin 1 (ZO-1, magenta) on the apical compartment of the transwell. The basolateral compartment containing the LMECs was stained for IAV NP (green) and the endothelial cell marker Vascular-Endothelial Cadherin (VE-CAD, magenta). In both cases the nuclei were visualized with Hoechst (scale bar 20 μm). (E) Epithelial cells (AO at ALI) or endothelial-epithelial co-cultures were inoculated with pH1N1, H1N1 or H3N2 virus at MOI 1. At 24 hours post-inoculation cytokines were measured in the apical compartment using the Legendplex assay. Data represented here show individual data points of cytokines derived from three independent experiments performed in biological duplicates and the mean +/- standard deviation (SD) is depicted. Mock of each condition was subtracted from the values of virus infected cells. Statistical significance was determined with Students-T-test (*<0.05, **<0.01, ***<0.005, ****<0.001).

**Figure 3.**
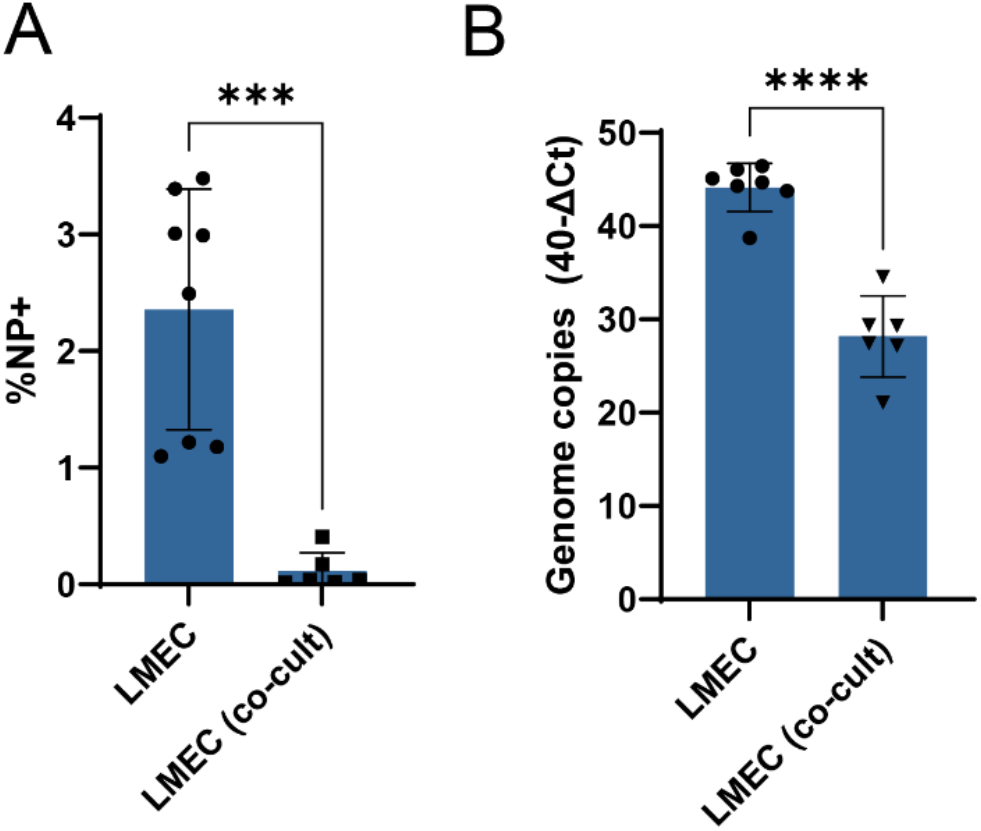
Quantification of influenza A virus infected cells in single endothelial cell cultures or in endothelial-epithelial co-cultures. Lung microvascular endothelial cells (LMECs) or differentiated airway organoids at air-liquid interface in co-culture with LMECs were inoculated with pH1N1 virus at MOI 1. (A) percentage of infection and (B) viral genome copies in LMEC single cultures compared to co-cultures were determined by flow cytometry or qRT-PCR at 24 hours post-inoculation. Data represented here show pooled data of virus titers derived from three independent experiments performed in biological duplicates and the mean +/- standard deviation is depicted. A student T-test was used to compare groups (*<0.05, **<0.01, ***<0.005, ****<0.001).

Next, we evaluated the pro-inflammatory cytokine response of infected epithelial cells in single culture or in co-culture in the apical compartment. We mainly detected pro-inflammatory cytokines (IL-6, IL-8, IP10) and IFNs (IFN-β, IFN-γ, IFN-λ1), in both single cultures and co-cultures. A more pronounced cytokine response was detected in the co-cultures compared to the single cultures at 24 hpi (Figure 2E). At 72 hrs no differences were observed between the different cultures (supplementary table 1). In conclusion, the AO at ALI were highly susceptible and permissive to seasonal and pandemic IAV and replication was associated with a robust proinflammatory cytokine response and breakdown of epithelial barrier integrity.

**Table 1.** Subtype and clade of 2019 viruses

| Virus | Subtype | Clade | Dutch identification number |
| --- | --- | --- | --- |
| H3N2 2019 | AH3N2 | 3C.3a | 19A01618 |
| H1N1 2019 | AH1N1pdm09 | 6B.1A5A | 19A01659 |

### Endothelial cells are not productively infected by influenza A viruses in epithelial-endothelial co-cultures but show an enhanced inflammatory response compared to single cultures

In subsequent experiments we focused on the LMECs in our epithelial-endothelial co-culture model. Even though all IAV robustly replicated in epithelial cells (Supplementary Figure 3A), we did not detect IAV NP in LMECs in these co-cultures at 24 or 72 hrs post epithelial cell inoculation (Figure 2B, D). This was in contrast to the LMECs in single culture where we found evidence for virus infection with immunofluorescence (Figure 1C-D). Next, we compared the infected LMECs in co-culture to LMEC single cultures and found significantly fewer infected cells in the co-cultures compared to infected LMEC single cultures (Figure 3A). In addition, significantly less viral RNA was detected in the LMECs of the co-cultures compared to inoculated LMECs in single cultures (Figure 3B). When LMECs were directly exposed to pH1N1, H1N1 or H3N2 virus by basolateral inoculation 24hrs after apical inoculation of epithelial cells, we rarely detected IAV NP in LMECs (Supplementary Figure 4).

**Figure 4.**
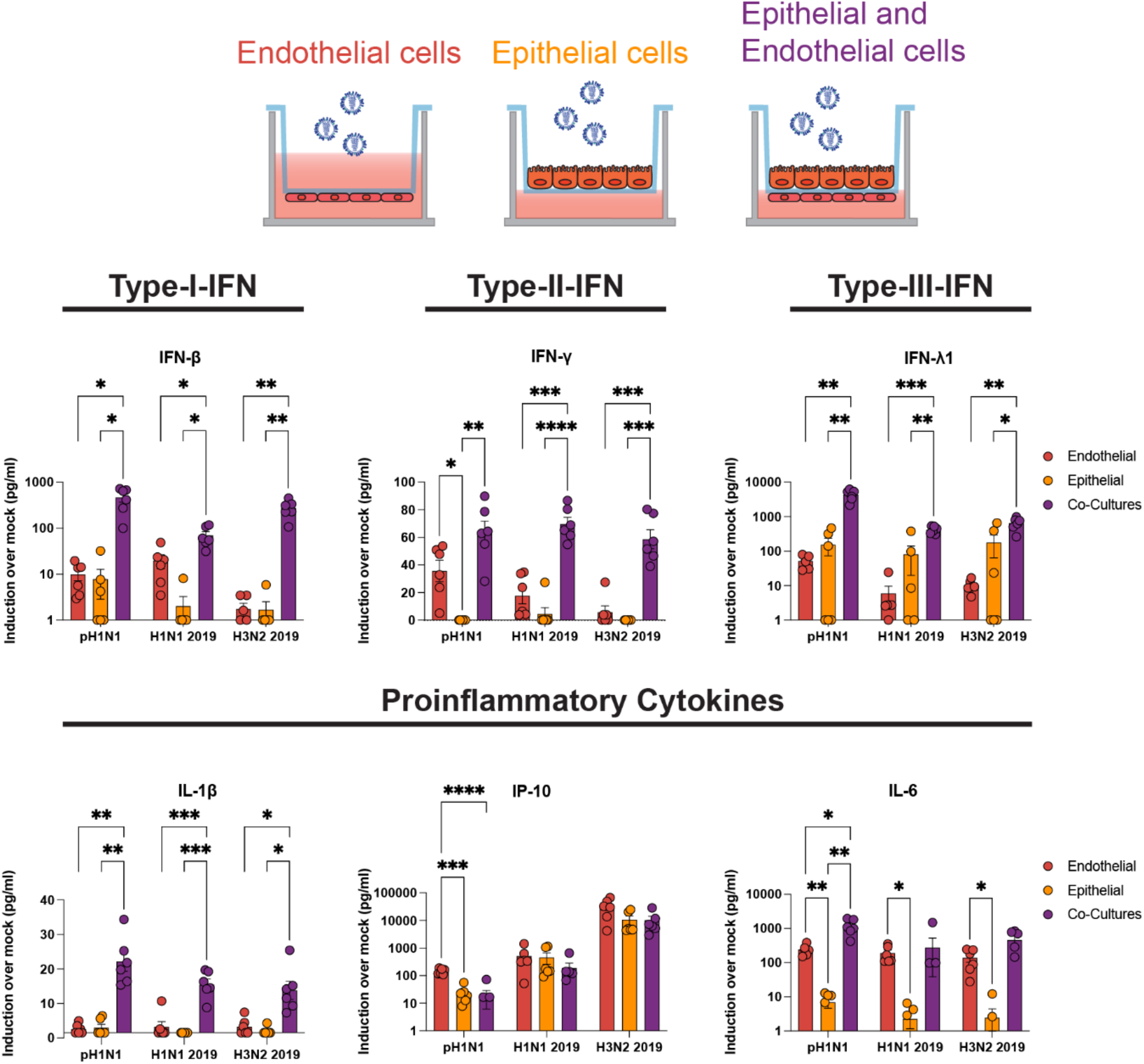
Cytokine production in endothelial cells in co-culture or in single culture. (A) Endothelial cells (lung microvascular endothelial cells), epithelial cells (airway organoids at air-liquid interface) and endothelial-epithelial co-cultures were inoculated with pH1N1, H1N1 and H3N2 virus at MOI 1. At 24 hours post-inoculation cytokines were measured in the basolateral compartment using a Legendplex assay. Data represented here show individual data points of cytokines derived from three independent experiments performed in biological duplicates and the mean +/- standard deviation is depicted. Mock of each condition was subtracted from the values of virus inoculated cultures. Statistical significance was determined with One-Way Anova and each group was compared to each other (*<0.05, **<0.01, ***<0.005, ****<0.001).

Next, we wanted to investigate the immune response of LMECs and AO at ALI in co-culture. No large differences in the upregulation of *IFIT1* or *IFN-β* gene expression were observed following pH1N1 inoculation. However, *IFN-λ* expression was significantly higher in the endothelial cells in endothelial cells compared to the epithelial cells. (Supplementary Figure 5). Finally, we investigated the cytokine secretion by endothelial cells in co-culture or in single cultures in the basolateral compartment (Supplementary table 1). Interferons (IFN-β, IFN-λ, IFN-γ) and pro-inflammatory cytokines (IL-1β, IP-10, IL-6) were upregulated upon virus infection in all cultures (Figure 4). Epithelial cells alone secreted mainly IFN-β and IFN-λ in the basolateral compartment but no IFN-γ, whereas endothelial cells alone were also capable of producing IFN-γ. Both epithelial cells and endothelial cells in single cultures produced IP-10 and IL-6 in the basolateral compartment, although endothelial cells seemed to contribute more to this production. Notably, we detected in the co-cultures higher concentrations of cytokines compared to the single cultures, which was more than the sum of the concentrations of cytokines produced in the single cultures. This effect was most pronounced for IL-1β but also observed for IFN-β, IFN-γ and IFN-λ. Other cytokines we measured were either above (IL-8) or below (IL-10) the limit of detection or showed no significant induction upon IAV infection (TNF-α, IL-12p70, IFN-λ2/3, GM-CSF, IFN-α2) (Supplementary table 1). There were no clear differences among the different viruses with respect to their cytokine profile.

**Figure 5.**
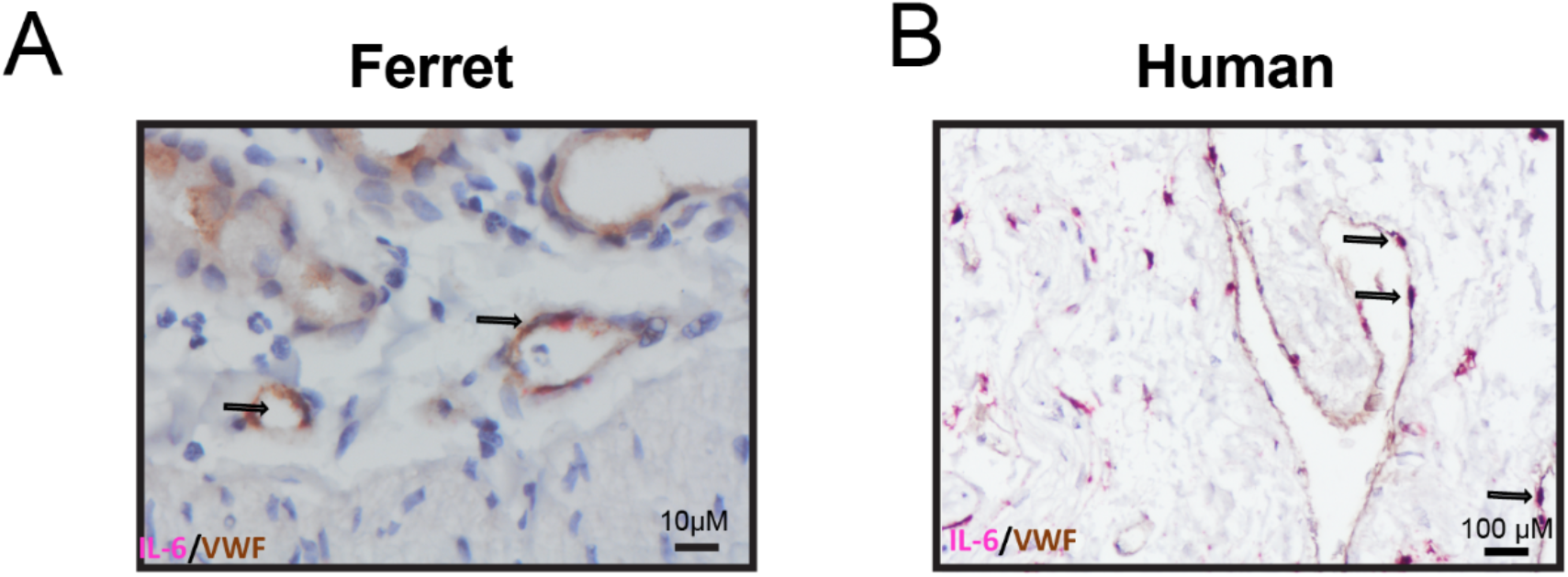
Endothelial cells in pH1N1-inoculated ferret and human lungs express IL-6 mRNA. (A) IL-6 production by endothelial cells was assessed in lung sections of a ferret inoculated with pH1N1 virus (1-day post-inoculation) and lung sections of human lung biopsies inoculated with pH1N1 virus (1-day post-inoculation) by In Situ Hybridization for IL-6, followed by immunohistochemistry using an antibody for endothelial cells (Von Willebrand Factor, VWF). Arrows indicate cells that are positive for IL-6 and VWF.

### Endothelial cells in *in vivo* pH1N1-infected ferret lungs and *ex vivo* pH1N1-infected human lungs express IL-6 mRNA

One of the main pro-inflammatory cytokines produced by LMECs in the co-culture model was IL-6. To confirm that endothelial cells contribute to the *in vivo* production of pro-inflammatory cytokines such as IL-6, we measured IL-6 mRNA in the lungs of ferrets inoculated with pH1N1 virus and in *ex vivo* human lung biopsies infected with pH1N1^22,23^. In both the ferret lungs and human lungs IL-6 mRNA transcripts were detected using *in situ* hybridization (ISH) in van Willebrand Factor (vWF) positive endothelial cells detected by immunohistochemistry (IHC, Figure 5A, B). This *in vivo* data supports the notion that endothelial cells contribute to IL-6 secretion.

## Discussion

In this study, we assessed the role of pulmonary endothelial cells in the pro-inflammatory responses upon IAV infection in a co-culture model consisting of AO at ALI and LMECs. We measured higher levels of pro-inflammatory cytokines in the basolateral compartment when endothelial cells were co-cultured with IAV-inoculated epithelial cells compared to infected endothelial or epithelial single cultures. This response was not associated with productive infection of the endothelial cells. One of the cytokines predominantly secreted by LMECs in the basolateral compartment was IL-6, which was also detected in endothelial cells in *vivo* in pH1N1 virus inoculated ferrets and in *ex vivo* inoculated human lung biopsies.

IAV infection of well-differentiated primary epithelial cells, either directly isolated from airway epithelial cells or stem cell-based, is well-characterized ^24,25^. Similar to our results, IAV was described to replicate efficiently in epithelial cells, cause damage to the epithelial barrier and induce robust amounts of pro-inflammatory cytokines (such as IL-1b, IL-6, IP-10, type I/III IFNs) in the apical compartment ^24,26–29^. Interestingly, here we show that when epithelial cells were co-cultured with endothelial cells the replication kinetics of all three IAV strains changed, and the levels of pro-inflammatory cytokines were increased. This suggests that the interaction between epithelial and endothelial cells in the respiratory tract influences the response to IAV infection.

The role of endothelial cells in the pathogenesis of influenza in humans is not well understood. Endothelial cells are rarely infected *in vivo* by zoonotic, pandemic and seasonal IAV, as measured by detection of viral antigen ^10,22,30,31^. Our study shows that small percentages of endothelial cells can be infected abortively by seasonal and pandemic influenza viruses when directly exposed to the viruses. In line with our data, other *in vitro* studies show low infection efficiency and abortive replication with seasonal or pandemic IAV ^11,14,16,21,22,31^. In contrast to the low percentage of infected endothelial cells measured by virus antigen, single cell RNA seq of PR8 (H1N1) infected mice lungs showed a high prevalence (∼50%) of viral RNA copies in pulmonary endothelial cells which correlated with high transcripts of ISGs ^18^. We also detected an induction of IFNs, ISGs and other cytokines in LMECs harvested from single cultures or co-cultures. This induction was most pronounced in co-cultures, in absence of detectable IAV antigen but presence of viral RNA. These data strengthen the hypothesis that abortive viral replication in LMECs induces an antiviral and inflammatory response likely through the detection of viral RNA by PRRs.

Several mechanisms can explain the induction of antiviral and inflammatory responses in endothelial cells when cultured together with infected epithelial cells. First, an abortive infection in LMECs, measured by qPCR, triggered an IFN response likely due to activating of PRRs. Our data shows that IAV breaks down the TER of AO at ALI and that endothelial cells can subsequently become abortively infected. Second, the trans-wells have pores (0.4 μm) which allow exposure of endothelial cells to cytokines produced by epithelial cells ^19^. Taken together, our results suggest that endothelial cells contribute to the inflammatory response to IAV infection by sensing viral RNA and/or epithelial-endothelial crosstalk and that both cell types are required when studying the context of IAV-induced inflammation in the lungs ^10,11,19^.

Endothelial cells line the blood vessels and cytokines produced by these cells will directly enter the bloodstream, potentially contributing to systemic inflammation in the blood. High levels of IL-6 and IFN-γ-induced IP-10 in the blood and bronchoalveolar lavage —cytokines that were produced to high levels by LMECS when co-cultured with epithelial cells—have been described as a biomarker for severe IAV disease in humans and mice ^32–36^. In addition, we showed that IL-6 mRNA is expressed by endothelial cells *in vivo* in ferrets and in *ex vivo* human lungs infected with pH1N1. As endothelial cells are prominent cells in the lung (30%)^18,37^, it is likely that endothelial cells are important contributors to the systemic inflammatory responses observed during severe influenza.

In conclusion, the described co-culture model suggests that abortively IAV-infected endothelial cells can fuel pro-inflammatory responses. Intervening or reducing the inflammatory responses in endothelial cells could potentially reduce morbidity and mortality associated with influenza.

## Material and Methods

### Primary Cells and Cell lines

Madin-Darby canine kidney (MDCK) cells were maintained in Eagle’s Minimal Essential Medium (EMEM; Lonza) supplemented with 10% FCS, 100 IU/ml penicillin, 100 μg/ml streptomycin, 2 mM glutamine, 1.5 mg/ml sodium bicarbonate (1 mM), 10 mM HEPES, and 0.1 mM nonessential amino acids. The medium was refreshed every 3 to 4 days, and cells were passaged at >90% confluence. Primary human lung microvascular endothelial cells (LMEC) were purchased at passage 3 from PromoCell-PromoKine (#C-12285) and cultured in 1% gelatine coated cell cultures flasks with Endothelial Cell Growth Medium MV-2 kit according to the manufacturer’s protocol (#C-22121, PromoCell-PromoKine), LMECs were only used up to passage 10 to ensure organ specificity. The cells were routinely checked for the presence of mycoplasma. Polyinosinic: polycytidylic acid (PolyI:C) was purchased from Invivogen and dissolved in water to a stock concentration of 1mg/ml.

### Viruses

Pandemic H1N1 virus (pH1N1 2009, A/Netherlands/602/2009) was isolated from a patient which visited Mexico. The isolate was propagated three times in MDCK cells. Seasonal H1N1 and H3N2 viruses were isolated in 2019 and were kindly provided by the National Influenza Centrum (Table 1). The received isolates were propagated once in MDCK cells.

### Human primary airway culture and differentiation

Human airway organoid cultures^38^ were generated and differentiated based on published protocols ^38,39^. To obtain differentiated organoid-derived cultures at air-liquid interface, organoids were disrupted into single-cell suspension with TrypLE Expressed and seeded on Transwell membranes (Corning) coated with rat tail collagen type I (Fisher Scientific) in airway organoid medium and complete base medium at a 1:1 ratio. When a monolayer was confluent (2-3 days), the apical medium was removed and the basolateral medium was replaced with complete base medium. The cultures were differentiated for 4-8 weeks with fresh medium every 5 days. Differentiation was visually confirmed by the production of mucus and ciliary movements or by antibody staining for tight junctions (zona occludens-1) and cilia (acetylated α-tubulin) and assessment by confocal laser scanning microscopy (Zeiss LSM700).

### Epithelial and Endothelial co-cultures and viral infections

The co-culture model of primary human epithelial and endothelial cells was established as follows. After full differentiation of human primary airway cultures on air liquid interface, the transwell inserts were inverted and coated with 150 µl 1% gelatin at 37°C for one hour. After the coating, the basolateral side of the transwell was washed once with PBS and 150.000 LMECs/well were seeded in 100µl MV2 medium in the basolateral compartment. To allow attachment of LMECs transwells were incubated at 37°C for one hour. Afterwards the basolateral compartment was filled with 750µl PneumaCult™-ALI Medium (#05001, Stemcell technologies, mixed with 750µl MV2 medium. For virus infection of the primary airway cultures at air liquid interface, first the apical cells were incubated with PBS containing Mg^2+^ and Ca^2+^ for 20 minutes and then the mucus was washed away by pipetting up and down. This process was repeated two times. According to the MOI, virus stocks were diluted in PBS containing Mg^2+^ and Ca^2+^ and 150µl of PBS and virus were added to the apical side. In case of epithelial single culture experiments the medium of basolateral compartment consisted of 1:1 mixture MV2 medium and Pneumacult, similar to the co-cultures. After 1 hour of attachment, the cells were washed three times with PBS containing mg2+ and ca2+. For infection of the endothelial cells, the transwell inserts were inverted and the LMECs were exposed to the corresponding MOI of virus diluted in MV2 medium. After one hour of incubation at 37°C, the basolateral side was washed three times with PBS and 750µl PneumaCult™-ALI Medium mixed with 750µl MV2 medium was added freshly.

### Trans-epithelial resistance measurements

A monolayer of epithelial cells grown on Transwell filters (0.4-μm-diameter pores, Corning, New York, NY) were infected with pH1N1, H1N1 and H3N2 virus with MOI 1 or left uninfected. The monolayer was analyzed for transepithelial resistance with an EVOM voltohmmeter (World Precision Instruments) with an STX-2 chopstick electrode at the indicated timepoint. The measured values were calculated by multiplying the electrical resistance by the area of the filter.

### Quantitative PCRs

Total RNA was isolated from cells using the High Pure RNA isolation kit (Roche) on a MagnaPure Machine and used for qRT-PCR. In short, 5 µl total RNA was amplified in a mix containing 5 µl of TaqMan Fast Virus 1-step master mix (Life Technologies), 1 µl primer-probe mix for β-actin, *IFN-β1, IFN-λ1, IFIT1* (Thermofisher Scientific) and 9 µl distilled water. The qRT-PCR temperature profile was 5 mins at 50 °C, 20 s at 95 °C, followed by 40 cycles of 3 s at 95 °C and 30 s at 60 °C. For determining the viral genome copies, 7 µl total RNA was amplified in a mix containing 5 µl of TaqMan Fast Virus 1-step master mix (Life Technologies), 0.4 µl primer-probe mix for IAV ^40^ and 9.6 µl distilled water. The qRT-PCR temperature profile was 5 mins at 50 °C, 20 s at 95 °C, followed by 45 cycles of 3 s at 95 °C and 30 s at 60 °C. The fold induction or genome copies fold change to input was calculated with the Double Delta Ct method^41^.

### Titration

Virus titers were determined by endpoint dilution on 30.000 MDCK cells/ well plated in 96-wells. 10-fold serial dilutions of cell supernatant in technical triplicate were prepared in infections medium consisting of EMEM supplemented with 100 IU/ml penicillin, 100 μg/ml streptomycin, 2 mM glutamine, 1.5 mg/ml sodium bicarbonate, 10 mM HEPES, 1× (0.1 mM) nonessential amino acids, and 1 μg/μl tosylsulfonyl phenylalanyl chloromethyl ketone (TPCK)-treated trypsin (Sigma-Aldrich). Before adding the titrated supernatants, the MDCKs were washed once with plain EMEM medium to remove residual FCS. 100 µl of the titrated supernatants were used to inoculate MDCKS and after 1-hour, infectious supernatant was removed and 200µl fresh infection medium was supplemented. Three days after infection, the supernatants of the MDCKs cells were tested for agglutination. For that 25µL of supernatant was mixed with 75µL 0.33% turkey red blood cells and incubated for 1 h at 4°C. Infection titers were calculated according to the method of Kärber and expressed as TCID50/ml^42^.

### Immunofluorescence Labeling

Cells on transwells were fixed by adding 4% formalin (1ml to the basolateral side and 500µl apical side) to the transwells for 30 minutes. Afterwards cells were washed three times with PBS and permeabilized with PBS containing 1% Triton X-100 followed by a 30 min incubation in blocking solution consisting of PBS supplemented 0.5% Triton X-100 and 1% bovine serum albumin (BSA) at room temperature. With a scalpel, the transwells were cut out of the plastic frame, the membrane was cut in half and washed three times in PBS. Primary antibodies concentrations were used according to the Table 2, were diluted in blocking solution and incubated at room temperature for one hour. Following a washing step where membranes were dipped three times in PBS, they were stained with the corresponding secondary antibodies, and Hoechst to visualize the nuclei, at room temperature for one hour. Stained membranes were afterwards washed three times in PBS and once in water and mounted in ProLong Antifade Mountant. Membranes were imaged using a Zeiss LSM 700 confocal microscope.

**Table 2.** Antibodies used for Immunofluorescence Microscopy.

| Antibodies | Cat nr | Final Concentration | Dilution | Manufacturer |
| --- | --- | --- | --- | --- |
| NP | EBS-I-047 | 2,0 µg/ml | 1:1000 | EVL laboratories |
| VE-CAD | AF938 | 2,0 µg/ml | 1:100 | R&D Systems |
| Acetylated-α-tubulin | SC-23950 | 2,0 µg/ml | 1:100 | Santa Cruz Biotechnology |
| ZO-1 | 339100 | 5,0 µg/ml | 1:100 | Life Technologies/InvitroGen |
| Hoechst | 62249 | 2 µM | 1:10.000 | Life Technologies/InvitroGen |

### Multiplexed bead assay for cytokine profiling

Cytokine concentration of the basolateral compartments were measured using the human antivirus response panel (13-plex) kit (LEGENDplex BioLegend). The kit was used according to the manufacturer’s manual. The data were analyzed with flow cytometry and final analysis of the concentration was performed using LEGENDplex analysis software v8.0.

### FACS stainings

LMECS and MDCKs were first washed with PBS and then released with trypsin and collected in a V-bottom plate. The cells were permeabilized using cytofix/cytoperm (BD Biosciences) for 20 minutes and blocked with 10% normal goat serum for 30 minutes. The cells were incubated at 4°C for 30 minutes with an unconjugated antibody for IAV NP (HB65, 8,0 μg/ml) followed by a PE-conjugated goat anti-mouse secondary antibody (DAKO) and measured on a flow cytometer (BD FACSLyric).

### Pathologic examination

Membranes with co-cultured epithelial and endothelial cells were collected and fixed in 10% neutral-buffered formalin for 30 min, cut in half and embedded in paraffin. For examination by light microscopy, 3 μm sections were deparaffinized and stained with hematoxylin and eosin (HE). Sections were also stained with periodic acid Schiff (PAS) for detection of mucoid cells according to standard methods.

### Influenza IHC

For detection of IAV nuclear protein (NP) in co-cultured epithelial and endothelial cells, 3 μm formalin-fixed, paraffin-embedded cross sections of the co-culture membranes were deparaffinized and rehydrated. NP antigen was retrieved by incubating sections briefly for 2 minutes in 0.1% protease at 37°C. The slides were then washed with phosphate-buffered saline (PBS)/0.05% Tween 20 and incubated with mouse IgG2a-anti-IAV NP (Clone Hb65, EVL laboratories) 5 μg/ml or mouse IgG2a isotype control (MAB003, R&D systems) 2.5 μg/ml in PBS/0.1% bovine serum albumin (BSA) for 1 hour at room temperature (RT). After washing, sections were incubated with horseradish peroxidase labeled goat-anti-mouse IgG2a (STAR133P, Serotec) 10 μg/ml in PBS/0.1% BSA for 1 hour at RT. Peroxidase activity was revealed by incubating slides in 3-amino-9-ethylcarbazole (AEC) (Sigma, St Louis, MO, USA) for 10 minutes, resulting in a bright red precipitate, followed by counterstaining with hematoxylin. A lung section from an experimentally influenza (pH1N1) inoculated ferret was used as a positive control.

### *In situ* analysis of ferret and human endothelial cells expressing IL-6

For determination of expression of IL-6 by endothelial cells upon influenza virus infection, lung sections of a ferret inoculated with pH1N1 virus (1 day post-inoculation (dpi)^22^ and human lung slices inoculated with pH1N1 (1dpi)^23^ were double stained by ISH for IL-6 using the RNAScope™ platform, followed by IHC using an antibody for endothelial cells (Von Willebrand Factor, VWF). RNA probes were designed by ACD Biotechne for ferret IL-6 (300031) and for human IL-6 (310371). First, ISH was performed on 3 μm formalin-fixed, paraffin-embedded lung sections using RNAScope™ Reagent Kit v2–RED (322350) as described by the manufacturer, up to amplification step 6. Slides were then washed in PBS/0.05% Tween 20 and incubated with rabbit anti human VWF (A008229-2, DAKO/Agilent tech) 1/500 or rabbit IgG isotype control (AB-105-C, R&D systems) 1/100 in PBS/0.1% BSA for 1 hour at RT. After washing, sections were incubated with biotinylated goat polyvalent anti-Mouse IgG (H+L) and anti-Rabbit IgG (H+L) (TP-125-HL, LabVision) for 10 min at RT, followed by horseradish peroxidase labeled streptavidin (D0397, DAKO/Agilent tech) 1/300 in PBS/0.1% BSA for 30 min at RT. Slides were washed and IL6 RNA molecules were visualized as red chromogenic dots according to the RNAScope™ protocol. Peroxidase activity from VWF IHC was revealed by incubating slides in 3,3’– diaminobenzidine-tetrachlorhydrate (DAB) (Sigma) for 3-5 minutes, resulting in a brown precipitate. Slides were counterstained with Gill’s hematoxylin, dried at 60C for 5-10 min and mounted with Ecomount.

### PolyI:C Treatment

Subconfluent LMECs were plated on 24-wells coated with 1% gelatine. On the next day, LMECs were transfected with 10 µg/ml or 1µg/ml PolyI:C with Lipofectamin 3000 (Invitrogen) according to the manufacturer’s protocol. After 24 hrs, cells were harvested in lysis buffer and RNA was extracted with the MagnaPure machine.

### Statistical Analysis

Statistical differences between experimental groups were determined as described in the figure legends. P values of ≤0.05 were considered significant. Graphs and statistical tests were made with GraphPad Prism version 9. Figures were prepared with Adobe Illustrator CC2019, Adobe Photoshop CC2019 and Biorender.

## Supporting information

Supplementary Part1

## Acknowledgement

D.V.R. is supported by fellowships from the Netherlands Organization for Scientific Research (VIDI contract 91718308) and a EUR fellowship.

## Contributions

Conceptualization L.B., L.R, D.V.R

Investigation L.B., L.R, L.L, F.F.W.B

Formal Analysis L.B., L.R, L.L, R.L.D.S, R.D.D.V, D.V.R

Resources L.B., L.R, L.L, D.N., M.M.L, B.M.H., R.L.D.S, R.D.D.V, D.V.R,

Supervision L.B, D.V.R

Visualization L.B, L.R

Writing Original Draft L.B., L.R, D.V.R

Writing-Reviewing all authors

Funding acquisition R.L.D.S, R.D.D.V, D.V.R

