## Supplementary Part1 for "The pro-inflammatory response to influenza A virus infection is fueled by endothelial cells"

**Bauer and Rijsbergen et al. 2022**

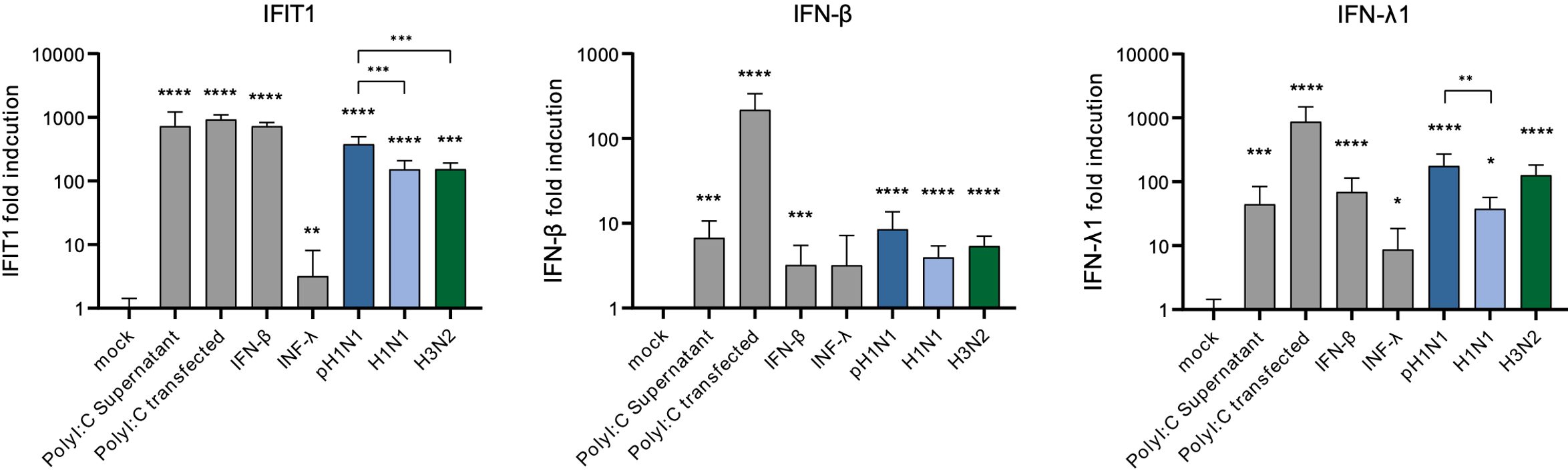

**Supplemental Figure 1.** **Interferon and interferon-stimulated gene expression in lung microvascular endothelial cells.** To evaluate the immunocompetence of lung microvascular endothelial cells (LMECs), the cells were exposed to recombinant IFN-β, IFN-λ, or the Toll-like receptor 3 agonist Polyinosinic:polycytidylic acid **(**Poly I:C) either in the supernatant or transfected. Alternatively, LMECs were inoculated with either pH1N1, H1N1 or H3N2. The gene expression of *IFN-β*, *IFN-λ* and interferon-stimulated gene *IFIT1* was analyzed via q-RT-PCR. Data represent mean +/- standard deviation (SD) from at least two independent experiments performed in biological triplicates. A student T-test was used to compare each group to mock and a one-way ANOVA was done to compare pH1N1, H1N1 H3N2 (*<0.05, **<0.01, ***<0.005, ****<0.001).

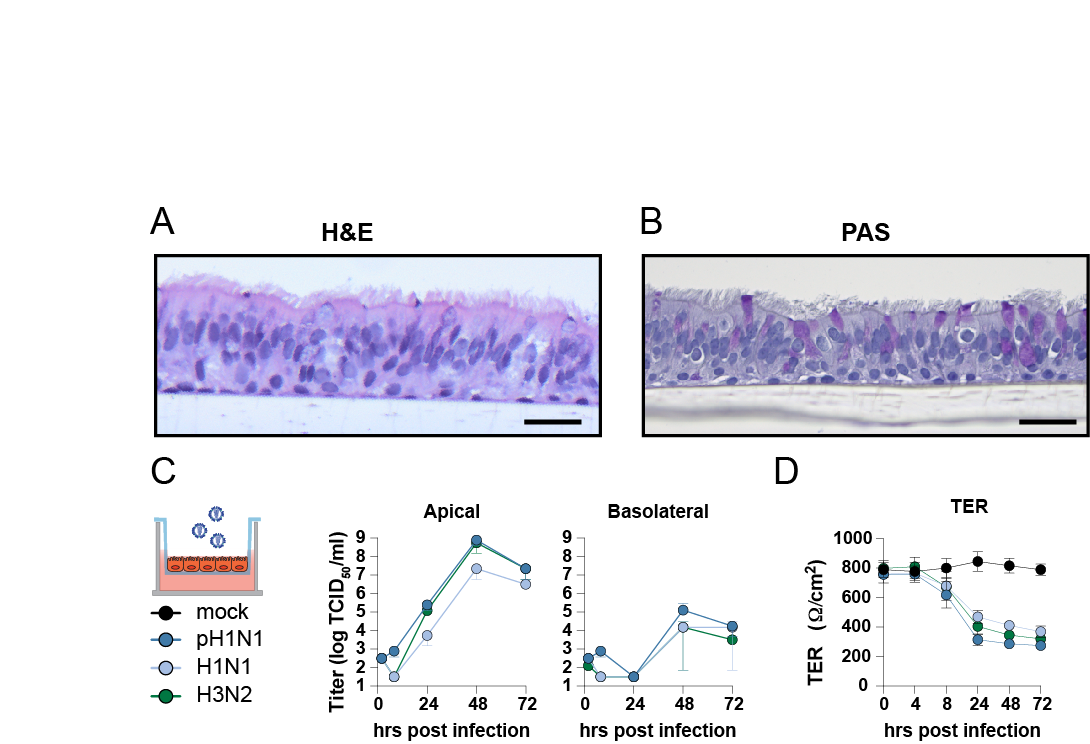

**Supplemental Figure 2. Influenza A virus replication in well-differentiated airways organoids at air-liquid interface.** Airway organoids were differentiated on the apical side of a transwell at air-liquid interface and tissue architecture, including presence of ciliated cells and goblet cells was confirmed with (A) H&E and a (B) PAS staining. (C) Growth kinetics of pH1N1, H1N1 and H3N2 in well-differentiated airways organoids using MOI 1. Virus titers were determined in the supernatants of the apical and basolateral side with endpoint dilution. (D) Determination of Trans-Epithelial resistance (TER) of influenza A virus-infected epithelial cultures at MOI 1 over time. Data are expressed relative to the TER value recorded prior to infection which was defined as 100%. Data represented here show pooled data of virus titers and TER values derived from three independent experiments performed in biological duplicates and the mean +/- standard deviation is depicted. Scale bar: 20 μm.

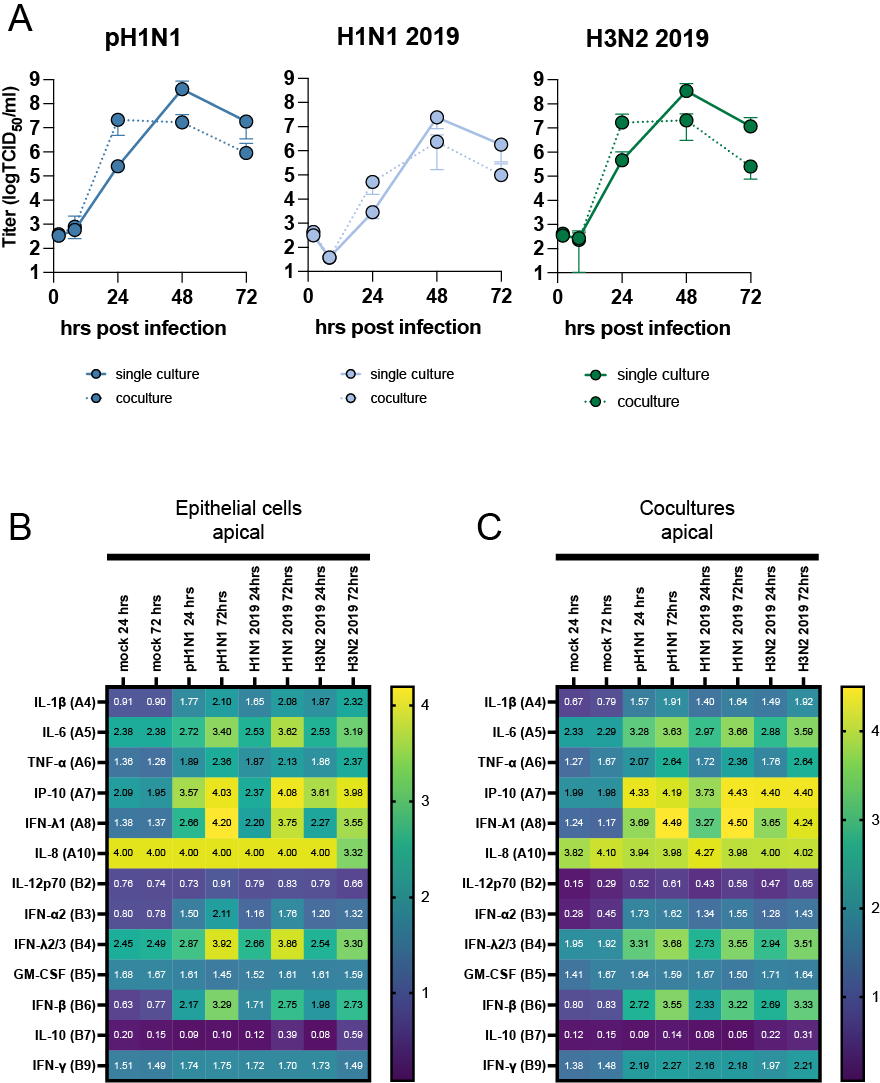

**Supplementary Figure 3**. **Influenza A virus replication in the apical compartment of epithelial-endothelial co-cultures.** (A) well differentiated airway organoids in co-cultures with and without lung microvascular endothelial cells were inoculated with pH1N1, H1N1 and H3N2 at MOI 1. At indicated timepoints virus titers were determined in the apical supernatants. Data represented here show log transformed pooled data of cytokines derived from three independent experiments performed in biological duplicates and the mean is depicted.
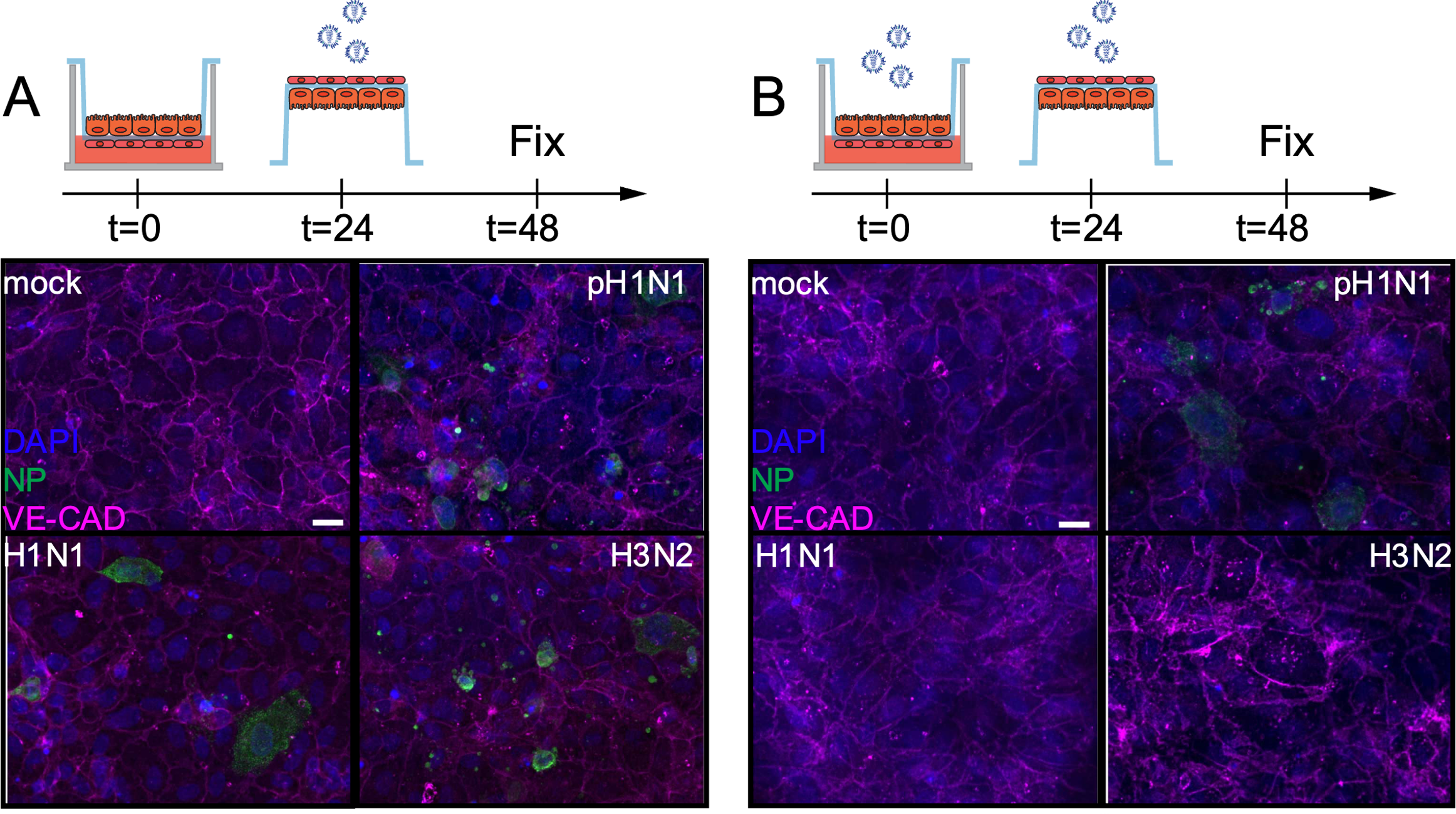

**Supplement Figure 4. No influenza A virus infected endothelial cells in epithelial-endothelial co-cultures when inoculated directly.**  (A) Well differentiated airway organoids in co-cultures with lung microvascular endothelial cells were inoculated with mock or (B) pH1N1, H1N1 and H3N2 at MOI 1. At 24 hours post-inoculation the endothelial cells were directly inoculated with homologous viruses. The cells were fixed after 24 hours post second inoculation and stained for the endothelial cell marker Vascular Endothelial-Cadherin (VE-CAD magenta) and influenza A virus nucleoprotein (NP, green). Hoechst (blue) was used to visualize nuclei. Scale bar: 20µM.

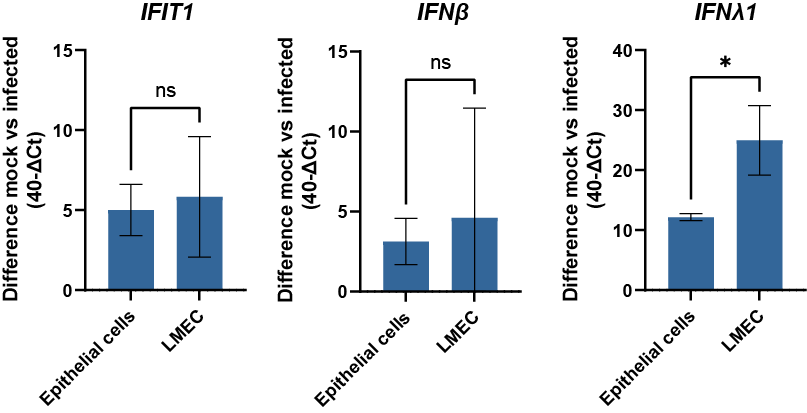

**Supplementary Figure 5.** **Interferon and interferon-stimulated gene expression in airway organoids at air-liquid interface and lung microvascular endothelial cells inoculated with pH1N1.** To evaluate the innate antiviral state of lung microvascular endothelial cells (LMECs) and epithelial cells (airway organoids at air-liquid interface) the cells were inoculated with pH1N1. The gene expression of interferon-stimulated gene *IFIT1*, IFN-βand IFN-λ was analysed via qRT-PCR. Data represent mean +/- standard deviation (SD) from at least two independent experiments performed in biological duplicates. A student T-test was used to compare each group to mock (*<0.05).

**(see Excel SHEET)**

**Supplementary table 1. Cytokine profiling in endothelial, epithelial and endothelial-epithelial co-cultures infected with pH1N1, H1N1 or H3N2**. Endothelial cells (lung microvascular endothelial cells), epithelial cells (airway organoids at air-liquid interface) and endothelial-epithelial co-cultures were inoculated with pH1N1, H1N1 and H3N2 at MOI 1. At 24 and 72 hours post-infection cytokines were measured in the apical and basolateral compartment using the human antiviral Legendplex assay (Biolegend). Raw data for all cytokines is represented here and is derived from three independent experiments performed in biological duplicates.
